# Impairing NK-mediated immune rejection through NKG2DL editing to improve CAR-T cell persistence

**DOI:** 10.64898/2026.04.07.716907

**Authors:** Hui Shi, Yi Wang, Xin Tang, Guangna Liu

**Author notes:** Correspondence to Guangna Liu.

## Abstract

CAR-T immunotherapy has achieved remarkable efficacy in hematologic malignancies. However, the widespread clinical adoption of autologous CAR-T products remains constrained by high costs, lengthy manufacturing process, and limited accessibility. Universal or off the shelf CAR-T (UCAR-T) cells derived from healthy donors offer a promising alternative, enabling immediate treatment at a lower cost. However, the allogeneic nature of UCAR-T cells triggers immune rejection by the host immune system after infusion, thereby compromising their persistence and therapeutic efficacy. Current strategies to circumvent this rejection focus on disrupting HLA class I expression. Although this modification allows UCAR-T cells to successfully evade T cell mediated elimination, the loss of HLA class I molecules renders them vulnerable to attack by host natural killer (NK) cells. In contrast to previous approaches that attempt to retain certain non-classical HLA molecules (such as HLA-E or HLA-G) to inhibit NK cells, we directly focused on editing the ligands that mediate NK cell rejection. Through transcriptomic and in vitro validation analyses, we found that UL16 binding proteins (ULBP) 2/5/6 were substantially upregulated in UCAR-T cells compared with nontransduced donor T cells. Elevated ULBP expression effectively activates the NKG2D receptor on allogeneic NK cells and leads to killing of UCAR-T cells, thereby impairing UCAR-T function. To test whether abrogating this NK activating signal could improve UCAR-T persistence and antitumor efficacy, we generated ULBP knockout UCAR-T cells using CRISPR-Cas9 editing. Deletion of ULBP2/5/6 significantly reduced NK cell mediated killing in vitro without affecting CAR expression or T cell effector function. Compared with wild type UCAR-T cells, ULBP deficient UCAR-T cells exhibited enhanced tumor killing efficacy in the presence of NK cells. Collectively, our findings identify ULBP upregulation as one of the mechanisms underlying NK cell mediated rejection of HLA deficient UCAR-T cells. Targeted ablation of ULBP molecules provides a novel strategy to confer resistance to host NK cells, thereby improving the therapeutic potential of universal CAR T products.

## Introduction

Chimeric antigen receptor T-cell therapy couples a single-chain variable fragment recognizing tumor-associated antigens with T-cell activation signals, thereby endowing T cells with the capacity for precise tumor killing ^[1]^. Although autologous CAR-T has demonstrated remarkable efficacy in hematologic malignancies^[2]^, its highly personalized manufacturing process presents several notable limitations, such as prolonged production times, complex procedures, high failure rates, and a cost of hundreds of thousands of dollars per single infusion, which severely restricts its accessibility^[3]^. To address these challenges, researchers have developed universal CAR-T. UCAR-T uses T cells from healthy donors as starting material and employs gene-editing technologies to disrupt the T-cell receptor^[4]^, thereby eliminating the risk of graft-versus-host disease and enabling large-scale industrial manufacturing to generate off-the-shelf cell products^[5, 6]^. However, when these allogeneic cells are infused into recipients with intact immune systems, they must not only evade host immune attack but also persist sufficiently to achieve complete tumor clearance, presenting UCAR-T with more complex challenges than those faced by autologous CAR-T^[7-10]^.

To address immune rejection issues by host immune system in UCAR-T therapy, current strategies typically involve disrupting the B2M and CIITA genes to eliminate human leukocyte antigen class I and class II expression, thereby preventing recognition by host CD8^+^ and CD4^+^ T cells and mitigating T cell-mediated rejection ^[11-13]^. Although this strategy enables evasion of adaptive immune surveillance, it triggers innate immune alarm. Natural killer cells^[14]^, as the first line of defense against aberrant cells, are regulated by the balance between inhibitory and activating receptors on their surface^[15]^. Among these, inhibitory receptors ^[16]^ (such as the KIR family and NKG2A) primarily recognize HLA class I molecules on target cells ^[17, 18]^. When NK cells encounter normal cells, HLA-I molecules engage inhibitory receptors, delivering inhibitory signals that prevent NK cell-mediated killing. Conversely, when NK cells encounter UCAR-T cells that have completely lost HLA-I expression, the inhibitory signals are lost, activating signals predominate^[19]^, and NK cells promptly initiate killing of UCAR-T cells^[20-22]^. Thus, for strategies that involve complete HLA I ablation (such as knockout of the B2M gene), how to deal with NK cell-mediated host-versus-graft rejection (HvGR) has become another critical issue.

Current research commonly attempts to retain partial HLA-I expression, for instance by selectively knocking out HLA-A and HLA-B^[23]^, or by overexpressing non-classical HLA molecules such as HLA-E/G on a B2M deficient background^[20, 24]^. However, leaked NK cell killing still exists even after these modifications. This study aims to develop a novel universal CAR-T cell platform devoid of HvGR risk, aimed at enhancing the in vivo persistence of UCAR-T cells and improving their anti-tumor efficacy.

## Methods

### Plasmid Construction

Plasmids were constructed via homologous recombination. Primers with 15-20 bp homologous sequences at their 5’ ends were synthesized by TSINGKE. PCR amplification was performed using KOD HiFi DNA Polymerase. The vector was digested with SalI and NotI restriction endonucleases (NEB). PCR products and digested fragments were separated by agarose gel electrophoresis, purified using the Tiangen Gel

Extraction Kit, and quantified by OD260 measurement. Ligation reactions were carried out with the ClonExpress II One-Step Cloning Kit (Vazyme). The ligation products were transformed into DH5α competent cells (Transgene) by incubation on ice for 30 min, heat shock at 42°C for 45 s, and recovery for 30-60 min, followed by plating on antibiotic-containing plates. Single colonies were screened by colony PCR, and positive clones were verified by sequencing. Plasmids were extracted using the Tiangen Midiprep Kit and quantified using a microspectrophotometer (MiuLab ND-100).

### Lentivirus Production and T cell Transduction

Lenti-X 293T cells were transfected with the target plasmid, pMD2.G, pRSV-Rev, and pMDLg at a mass ratio of 4:1:1:2 using PEI as the transfection reagent. The culture medium was replaced 14-16 hours post-transfection, and viral supernatants were collected at 48 and 72 hours, followed by concentration using PEG8000. Viral titers were determined based on the percentage of RFP-positive Jurkat cells at 72 hours post-transduction. For human T cell transduction, 5×10^5^ T cells were mixed with concentrated virus at a multiplicity of infection (MOI) of 3-10 and cultured for 24 hours prior to assessment of transduction efficiency.

### Construction of universal CAR-T cells

To activate human primary T cells, CD3 antibody (Biolegend) was pre-coated overnight at 4°C. Thawed PBMCs (Bokang Bioengineering) were centrifuged at 1500 rpm and seeded into pre-coated plates at 1×10^6^ cells/mL in RPMI 1640 medium (Thermo Fisher). After 24 hours of activation at 37°C, cells were transduced with CAR-carrying lentiviral particles. Twenty-four hours post-transduction, 100 pmol of sgRNA (GenScript) and 31 pmol of Cas9 protein (Kactusbio) were added per 1 × 10^6^ cells in the reaction system, and nucleofection was performed using the P3 Primary Cell 4D-Nucleofector™ X Kit S (LONZA) to knock out B2M, *TRAC*, and other genes. Subsequently, cells were maintained at a density of 1-2 × 10^6^ cells/mL for further expansion.

### NK Cell Culture

To activate NK cells from PBMCs, CD2/NKP46 antibody (Thermo Fisher) was pre-coated overnight at 4°C. Thawed PBMCs (Bokang Bioengineering) were centrifuged at 1500rpm and seeded into the pre-coated plates at a density of 2 × 10^6^ cells/mL in OptiVitro NK Cell Medium P01 (Excellbio). After enrichment culture, cells were collected by centrifugation on day 7, followed by magnetic bead separation using CD3 Nanobeads (CytoSinct) with MACS LD columns (Miltenyi). NK cells were collected from the flow-through fraction and further cultured at a density of 1.5 × 10^6^cells/mL.Prior to the experiment, cells were stained with the flow cytometry antibodies CD3-PE-Cy7 and CD56-APC (Biolegend), and NK cell purity and viability were assessed using flow cytometry (BD).

### NK Cell Cytotoxicity Assay

NK cells and CAR-T cells were washed with 1×PBS, resuspended in staining buffer containing antibodies, incubated, washed twice, and analyzed using a BD flow cytometer. T cells were labeled with CFSE (Beyotime), and NK cells were co-cultured with CAR-T cells at the indicated effector-to-target (E:T) ratios. After 24 hours of co-culture at 37°C, cells were stained with Fixable Viability Dye eFluor ™ 450 for viability assessment and subsequently analyzed by BD flow cytometry. Calculation of cytotoxicity efficiency: Each group was used as a control with non-NK cell-embedded NE, i.e.: (Cell death rate in co-culture with NK cells-Cell death rate in groups without NK cells) = NK cytotoxicity rate against T cells.

### UCAR-T Cell Cytotoxicity Assay

Target tumor cells (Huh7-luc/GFP) were plated in cell-culture plate at a density of 8×10^5^ cells/mL, after 12 hours, 8×10^5^/ 4×10^5^ CAR-T cells/ well with or without 2.67×10^5^/ 4×10^5^ NK cells/ well were added to the plate. Then cells were co-cultured for an additional 24 hours at 37°C. Analysis was then performed using a luciferase assay kit (Yeason) with an Agilent multimode reader. The specific cytotoxicity rate was calculated using the following formula: Specific cytotoxicity rate (%) = [1-(experimental group RLU value -background control RLU value) / (maximum surviving control RLU value-background control RLU value)] × 100% (RLU: Relative Light Units).

### Data Analysis

Data analysis was performed using the following software: SnapGene 6.1.1 was used for primer design and plasmid sequence alignment; FlowJo 10.8.1 was used for flow cytometry data analysis and visualization; and GraphPad Prism 10.0.2 was used for cytotoxicity assessment and cytokine quantification. Statistical comparisons between groups were evaluated using two-way ANOVA. Statistical significance is denoted as follows: ^*^ means p<0.05, ^**^ means p<0.01, ^***^ means p<0.001, ^****^ means p<0.0001, and ns means no significant difference.

## Results

### NK cell-mediated rejection of UCAR-T cells profoundly impairs the effector functions of UCAR-T

To assess the cytotoxicity of NK cells on HLA class I-deficient UCAR-T cells, we utilized GPC3-targeted CAR-T cells and hepatocellular carcinoma model to carry out our study. To generate universal CAR-T cells, GPC3 CAR were transduced into primary human T cells, and B2M and TRAC genes were knocked out through CRISPR/Cas9 system (Figure 1A). Flow cytometry analysis revealed that UCAR-T cells in the experimental group successfully knocked out both the B2M and TRAC genes while maintaining successful expression of the CAR molecule (Figure 1B).When co-cultured with Huh7 tumor cells, UCAR-T cells exhibited comparable cytotoxicity to that of Mock CAR-T cells (Figure 1C). And as expected, due to the deficiency of HLA class I molecules, NK cells exerted robust cytotoxicity against UCAR-T cells which was significantly higher than that against Mock CAR-T cells (Figure 1D). Furthermore, to evaluate the impact of NK-mediated cytotoxicity on CAR-T cell function, we co-cultured Mock CAR-T or UCAR-T cells with tumor cells in the presence of NK cells (Figure 1E). Compared to Mock CAR-T cells, the effector function of UCAR-T cells against target cells was markedly impaired by NK cells (Figure 1F). Together, these findings indicate that NK cell-mediated rejection severely compromises the antitumor activity of UCAR-T cells.

**Figure 1.**
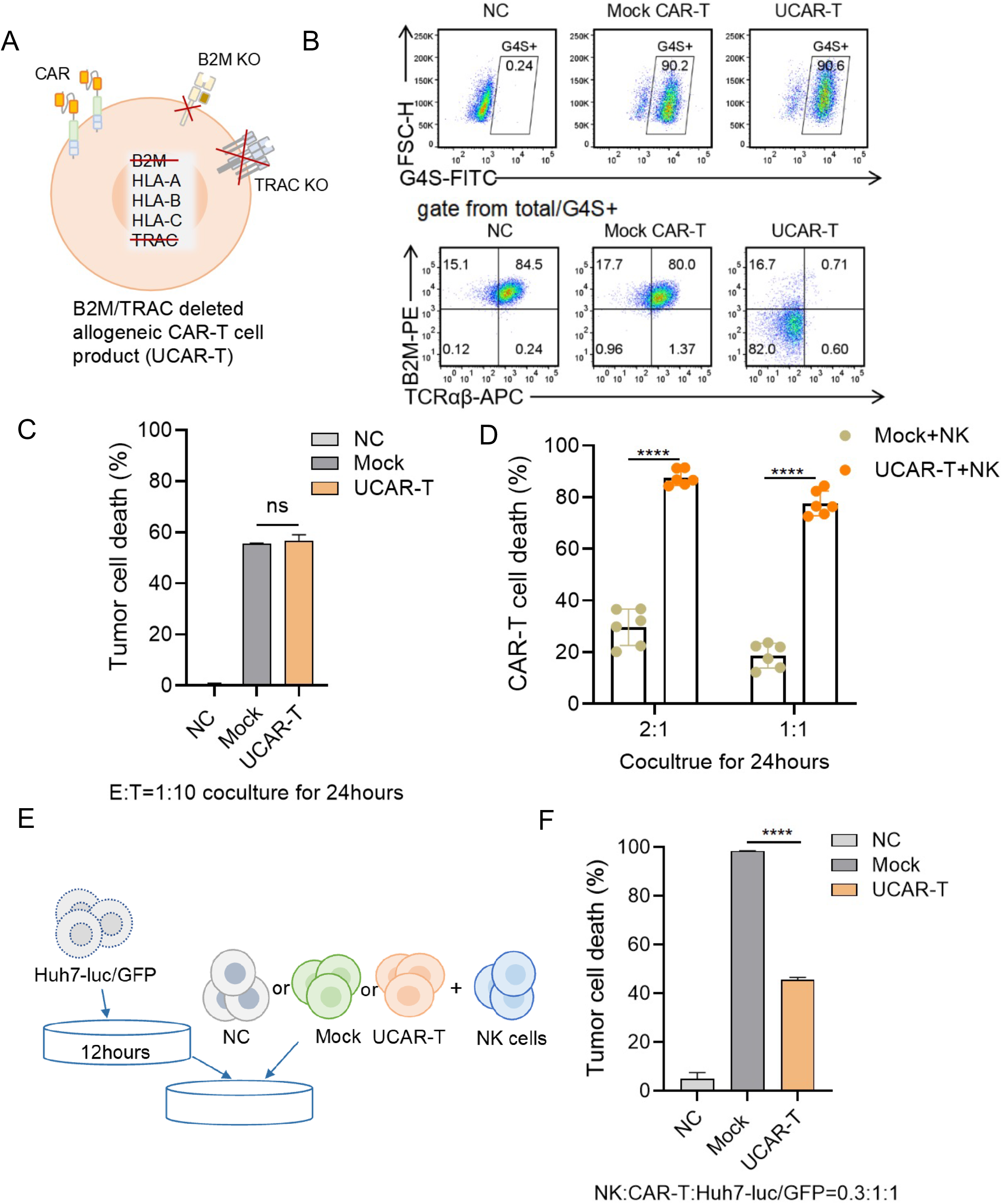
Evaluation of NK Cell-Mediated Rejection of UCAR-T. A. Schematic diagram of gene editing during UCAR-T cell generation: on Day 1 after PBMC thawing, cells were transduced with GPC3 CAR lentivirus; on Day 2, nucleofection was performed to knock out the B2M and TRAC genes in CAR-T cells B. Transduction efficiency and knockout efficiency of UCAR-T assessed by flow cytometry on Day 6. C. Killing efficiency of NC, Mock, and UCAR-T co-cultured with Huh7-luc/GFP target cells at an E:T ratio of 1:10 for 24 hours D. Detection after co-culture of NK cells (Day 8) with UCAR-T and Mock (Day 7) at E:T ratios of 2:1 and 1:1 for 24 hours. E. Schematic diagram of co-culture of NK cells with NC, Mock, UCAR-T, and Huh7-luc/GFP target cells F. Killing efficiency after NK cells were co-cultured with NC, Mock, or UCAR-T at an E:T ratio of 0.3:1 for 24 hours, followed by secondary co-culture with Huh7-luc/GFP target cells at a ratio of 0.3:1:1 for an additional 24 hours.Data represent the means ± SD of three technical replicates, statistical significance was determined by two-way ANOVA. ^****^ means p*<*0.0001 and ns means no significant difference.

### ULBP2/5/6 are the key targets mediating NK cell-mediated rejection of UCAR-T cells

To elucidate the mechanisms underlying NK cell-mediated rejection of UCAR-T cells, potential ligands involved in NK cell recognition were investigated. Transcriptome analysis revealed significant enrichment of NK cell-mediated cytotoxicity pathways in CAR-T cells compared with negative controls (NC) (Figure 2A). Among differentially expressed genes, several stimulatory NK receptors, such as ULBP1, ULBP2, and NCR1 were upregulated, suggesting activation of NK cell recognition-related pathways (Figure 2B). NK cells bind to target cells through activating receptors or adhesion molecule receptors, which recognize stress -induced ligands or adhesion molecules that are frequently overexpressed on tumor cells. (Figure 2C). Therefore, we next examined the activating receptors and key adhesion molecules of NK cells. Flow cytometry further confirmed substantial upregulation of ULBP2/5/6 on CAR-T cells, whereas other NK ligands showed minimal changes (Figure 2D). Notably, the adhesion molecules CD54 and CD58 were highly expressed on both CAR-T cells and NC cells (Figure 2E), implicating ULBP2/5/6 as key mediators of NK cell rejection.

**Figure 2.**
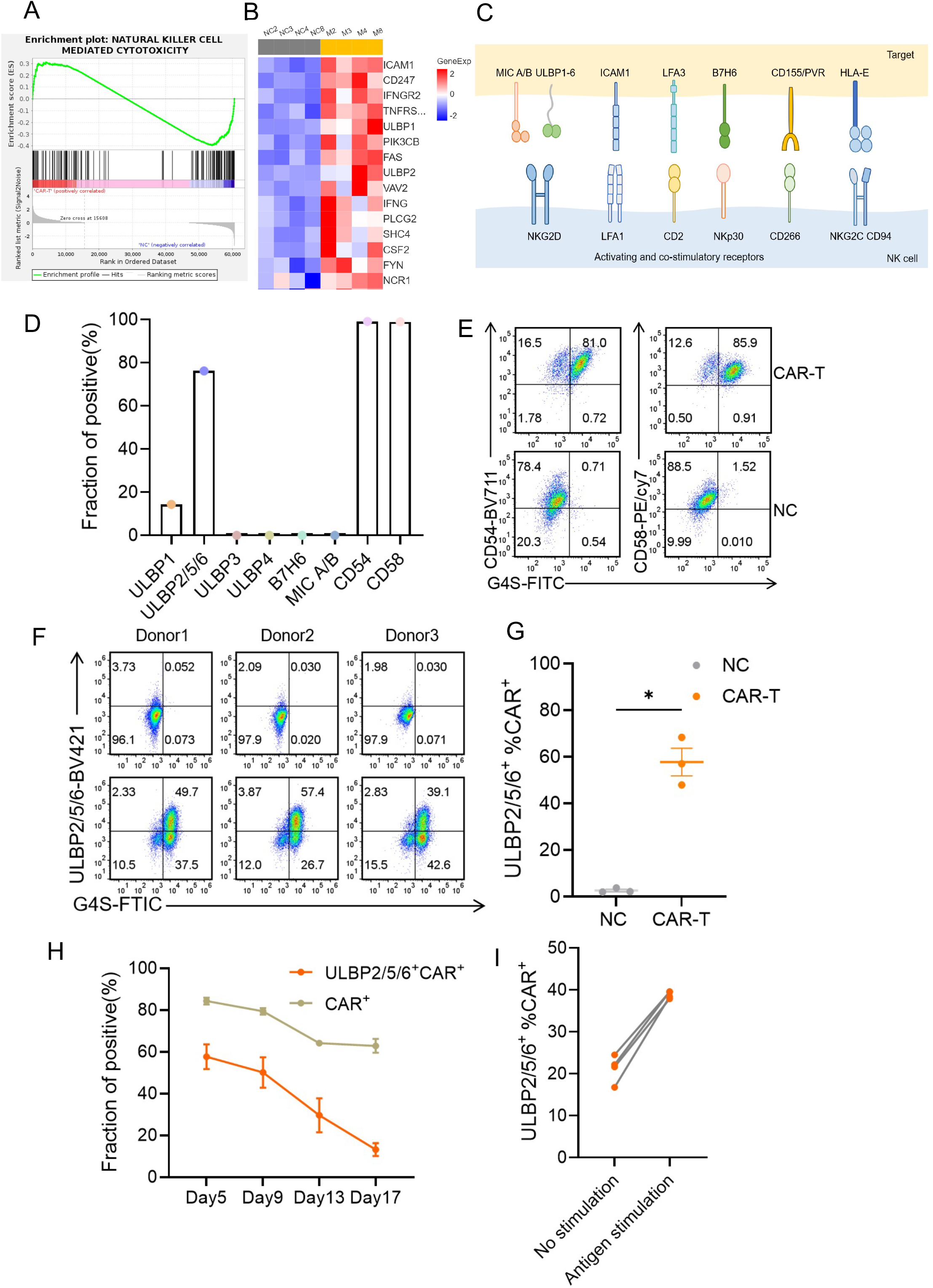
Expression of NK Cell Ligands on CAR-T Cells. A. Enrichment analysis of NK cell-mediated cytotoxicity based on transcriptome sequencing of CAR-T and NC cells collected from four different donors on Day 7 B. Heatmap of NK cell-mediated cytotoxicity-related gene expression based on transcriptome sequencing of CAR-T and NC cells collected from four different donors on Day 7 C. Schematic diagram of activating receptors and their ligands on NK cells D. Expression of NK cell ligands on CAR-T cells detected by flow cytometry E. Expression of CD54,CD58 on CAR-T/NC cells detected by flow cytometry F. Expression of ULBP2/5/6 detected by flow cytometry on Day 5 after GPC3 CAR transduction G. Statistical analysis of ULBP2/5/6^+^ CAR^+^ expression in NC and CAR-T cells on Day 5.Data represent the median of three technical replicates, statistical significance was determined by Paired t test, ^*^ means p*<*0.05. H. Longitudinal expression of ULBP2/5/6 ^+^ CAR ^+^ in relation to CAR ^+^ expression and culture duration, assessed by flow cytometry from Day 5 to Day 17. I. On Day 13, CAR-T cells were restimulated with GPC3 protein, and the expression of ULBP2/5/6^+^ CAR^+^ was examined.

Consistent upregulation of ULBP2/5/6 following GPC3 CAR transduction was observed across multiple donors (Figure 2F, G). Longitudinal analysis showed that the proportion of ULBP2/5/6^+^ CAR^+^ cells declined in parallel with CAR^+^ cells over time (Figure 2H), and restimulation with GPC3 protein increased ULBP2/5/6 expression (Figure 2I). Collectively, these results indicate that expression of ULBP2/5/6 is correlative with the expression of CAR and can be upregulated upon antigen stimulation.

### Disruption of ULBP2/5/6 function reduces NK cell-mediated rejection of UCAR-T cells

Given that ULBP2/5/6 molecules are ligands for NKG2D on NK cells, to investigate the contribution of NKG2D to NK cell-mediated rejection of UCAR-T cells, we added NKG2Dα to the co-culture system. This treatment significantly reduced NK cell-mediated killing of UCAR-T cells (Figure 3A, B), indicating that NKG2D and its ligands ULBP2/5/6 are key mediators of the rejection response. To further validate this mechanism, we knocked out the ULBP2/5/6 genes in UCAR-T cells to generate U-UCAR-T cells (Figure 3C). We next assessed the phenotypic characteristics, CAR expression, and anti-tumor function of U-UCAR-T. The results showed that the knockout procedure did not affect CAR expression or cytotoxic function (Figure 3D, E), thereby providing functionally intact cells for subsequent evaluation. Subsequently, we established a co-culture system of NK cells with U-UCAR-T to compare the ability of U-UCAR-T and conventional UCAR-T to resist NK cell cytotoxicity. When co-cultured with NK cells, U-UCAR-T cells exhibited enhanced resistance to NK cell cytotoxicity (Figure 3F).

**Figure 3.**
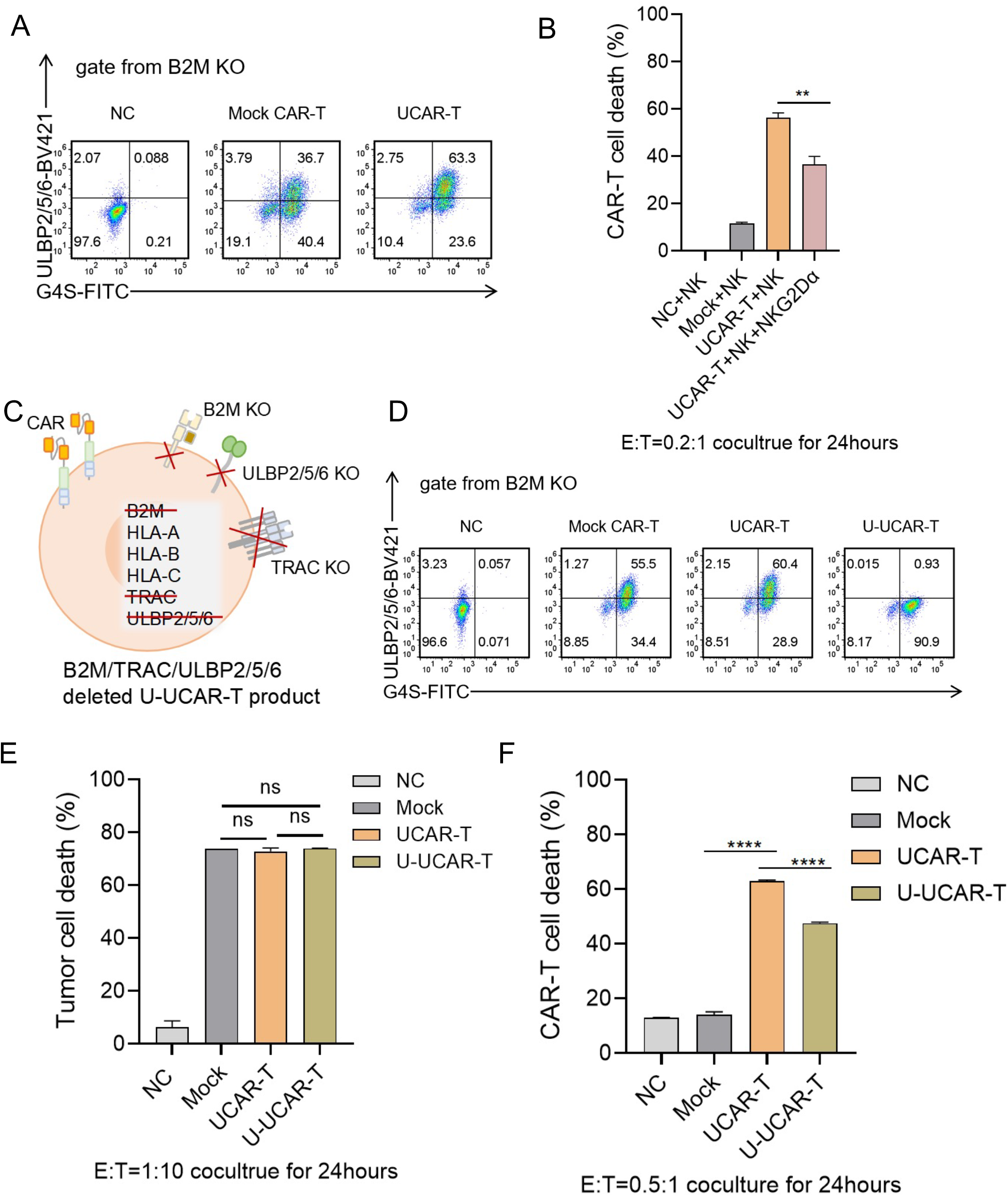
Evaluation of NK Cell-Mediated Rejection of U-UCAR-T. A. On Day 1, cells were transduced with GPC3 CAR; on Day 2, nucleofection was performed to generate UCAR-T. On Day 5, CAR membrane expression and ULBP2/5/6 expression were assessed by flow cytometry. B. On Day 8, NK cells were co-cultured with NC, CAR-T, UCAR-T, or UCAR-T plus NKG2D α (at a working concentration of 5 μg/mL) at an E:T ratio of 0.2:1. Killing efficiency was evaluated after 24 hours of co-culture. C. Schematic diagram of gene knockout for U-UCAR-T construction. D. On Day 1, cells were transduced with GPC3 CAR; on Day 2, nucleofection was performed to knock out B2M, TRAC, and ULBP2/5/6 in CAR-T cells. On Day 5, CAR membrane expression and ULBP2/5/6 expression were detected by flow cytometry. E. On Day 7, T cells were co-cultured with Huh7-luc/GFP target cells at an E:T ratio of 1:10, and killing efficiency was assessed after 24 hours. F. On Day 8, NK cells were co-cultured with NC, CAR-T, UCAR-T, or U-UCAR-T at an E:T ratio of 0.5:1, and killing efficiency was evaluated after 24 hours.Expression of ULBP2/5/6 detected by flow cytometry on Day 5 after GPC3 CAR transduction.Data represent the means ± SD of three technical replicates, statistical significance was determined by two-way ANOVA. ^**^ means p*<*0.01, ^****^ means p*<*0.0001 and ns means no significant difference.

### Abrogating ULBP2/5/6 maintains UCAR-T cell effector function in the setting of NK cell-mediated rejection

To comprehensively compare the ability of U-UCAR-T and conventional UCAR-T to maintain anti-tumor function in the presence of NK cells, we established a three-cell co-culture system containing NK cells, CAR-T cells, and tumor cells at various effector-to-target (E:T) ratios. Unlike conventional UCAR-T cells, which exhibited significantly impaired effector function under the same conditions, U-UCAR-T cells retained their cytotoxic function under NK cell-mediated rejection pressure (Figure 4A, B). In addition, we preliminarily explored whether ULBP2/5/6 knockout affects other important biological characteristics of UCAR-T, such as cytokine secretion. Consistent with the cytotoxicity results, conventional UCAR-T cells showed markedly reduced secretion of key pro-inflammatory cytokines (i.e., IL-2, IFN-γ, and TNF-α) after co-culture with NK cells. In contrast, U-UCAR-T cells exhibited significantly restored secretion levels of these cytokines under the same co-culture conditions (Figure 4C, D, E), further confirming at the functional level that U-UCAR-T maintains a favorable activation state and effector potential in the presence of NK cells. Collectively, these findings demonstrate that ULBP2/5/6 knockout not only enables UCAR-T to evade direct NK cell killing but, more importantly, preserves its ability to exert anti-tumor functions under inflammatory conditions.

**Figure 4.**
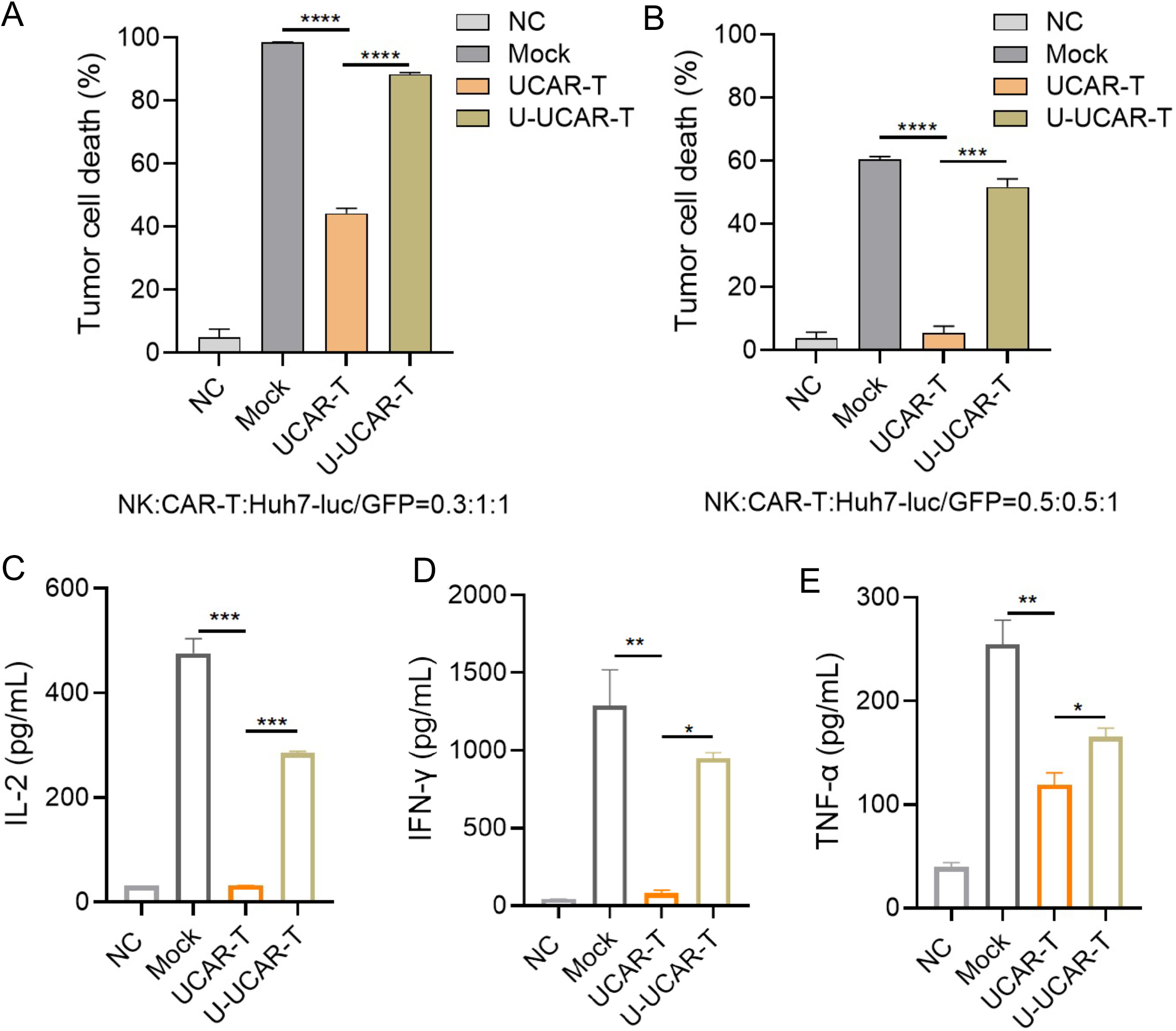
U-UCAR-T Exhibits Robust Effector Function under NK Cell-Mediated Rejection Pressure. A. Cytotoxicity after NK cells were co-cultured with NC, Mock, UCAR-T, or U-UCAR-T at an E:T ratio of 0.3:1 for 24 hours, followed by secondary co-culture with Huh7-luc/GFP target cells at a ratio of 0.3:1:1 for an additional 24 hours B. Cytotoxicity after NK cells were co-cultured with NC, Mock, UCAR-T, or U-UCAR-T at an E:T ratio of 0.5:1 for 24 hours, followed by secondary co-culture with Huh7-luc/GFP target cells at a ratio of 0.5:1:1 for an additional 24 hours C. Detection of IL-2 by ELISA after 24 hours of co-culture at an E:T ratio of 0.5:0.5:1 D. Detection of IFN-γ by ELISA after 24 hours of co-culture at an E:T ratio of 0.5:0.5:1 E. Detection of TNF-α by ELISA after 24 hours of co-culture at an E:T ratio of 0.5:0.5:1.Data represent the means ± SD of three technical replicates, statistical significance was determined by two-way ANOVA, ^*^ means p<0.05, ^**^ means p*<*0.01, ^***^ means p*<*0.001, ^****^ means p*<*0.0001.

## Discussion

This study overcame the technical bottleneck of NK cell in vitro expansion and established an efficient NK cell culture system, providing critical effector cells for rejection research. On this basis, we found that NK cells demonstrated robust cytotoxicity against UCAR-T cells engineered to eliminate HLA class I expression via B2M knockout, and this rejection severely compromised the subsequent anti-tumor function of UCAR-T. Through transcriptome sequencing and flow cytometric validation, we identified that ULBP2/5/6-major activating ligands for NKG2D-were significantly upregulated on the surface of UCAR-T cells, with their expression levels positively correlated with CAR expression and T-cell activation status. Functional blocking experiments confirmed that the addition of soluble NKG2Dα significantly reduced NK cell-mediated killing of UCAR-T, thereby revealing the NKG2D/ULBP2/5/6 axis as the core molecular mechanism driving NK cell rejection of UCAR-T.

Based on these findings, we employed CRISPR/Cas9 technology to further knock out ULBP2/5/6 in conventional UCAR-T (with B2M and TRAC double knockout), successfully generating triple-knockout U -UCAR-T cells. These cells maintained a high CAR-positive rate (>90%) while ULBP2/5/6 expression was effectively eliminated to below 1%, and the in vitro killing function against hepatocellular carcinoma cells remained comparable to that of conventional CAR-T. In vitro co-culture experiments demonstrated that U-UCAR-T exhibited significantly enhanced resistance to direct NK cell killing. In a three-cell co-culture system simulating host immune pressure (NK cells-CAR-T cells-tumor cells), U-UCAR-T sustained higher tumor-killing capacity following NK cell challenge, with secretion levels of effector cytokines (IL-2, IFN-γ, TNF-α) significantly restored compared with conventional UCAR-T. These findings confirm that ULBP2/5/6 knockout effectively protects UCAR-T from in vitro NK cell rejection without compromising their fundamental functions.

Although promising in vitro results were obtained in this study, it is challenging for an in vitro system alone to fully recapitulate the complex in vivo microenvironment that influences cell fate. In vivo, UCAR-T cells are subject not only to continuous immune surveillance by host NK cells, but also to multiple constraints imposed by the tumor microenvironment (TME). Moreover, critical processes such as distribution, homing, proliferation, persistence, and long-term functional maintenance of cells necessitate evaluation under conditions that more closely mirror the physiological context. Therefore, validating in vitro findings through in vivo studies is essential for assessing the clinical translational potential of U-UCAR-T cells. Therefore, the performance of U-UCAR-T in a relatively intact immune environment remains to be validated using more physiologically relevant animal models^[25]^, such as humanized PBMC-reconstituted mice^[26]^. We have not yet conducted in-depth assessments of the long-term persistence, tissue distribution, or potential long-term safety of U-UCAR-T in vivo. Moreover, simultaneous knockout of three genes may pose a risk of genomic instability^[27]^, making off-target analysis^[28]^, karyotyping, and long-term GVHD monitoring essential for subsequent studies. Additionally, whether the function of U-UCAR-T within the solid tumor microenvironment is affected by other immunosuppressive mechanisms (e.g., PD-1/PD-L1 pathway, myeloid-derived suppressor cells) ^[29-31]^and whether combination with other immunotherapeutic strategies could yield synergistic effects represent avenues worthy of further exploration^[32]^.

In summary, further knockout of ULBP2/5/6 in conventional UCAR-T (with B2M and TRAC double knockout) effectively blocks recognition and killing by NK cells via the NKG2D receptor, thereby generating immune-privileged UCAR-T. This optimized strategy significantly enhances the ability of UCAR-T to resist NK cell rejection both in vitro, while preserving normal anti-tumor effector function. Starting from a clinically relevant challenge, this study, through systematic mechanistic investigation and functional validation, provides a new theoretical basis and an effective solution to overcome host immune rejection-a major bottleneck in the clinical application of UCAR-T. These findings hold significant scientific importance and translational value for advancing the development of universal CAR-T cell therapy.

## Reference

[1] June CH, Sadelain M. Chimeric Antigen Receptor Therapy [J]. N Engl J Med, 2018, 379(1): 64–73.

[2] Maude SL, Laetsch TW, Buechner J, et al. Tisagenlecleucel in Children and Young Adults with B-Cell Lymphoblastic Leukemia [J]. N Engl J Med, 2018, 378(5): 439–48.

[3] Sterner RC, Sterner RM. CAR-T cell therapy: current limitations and potential strategies [J]. Blood Cancer J, 2021, 11(4): 69.

[4] Eyquem J, Mansilla-Soto J, Giavridis T, et al. Targeting a CAR to the TRAC locus with CRISPR/Cas9 enhances tumour rejection [J]. Nature, 2017, 543(7643): 113–7.

[5] Depil S, Duchateau P, Grupp S A, et al. ‘Off-the-shelf’ allogeneic CAR T cells: development and challenges [J]. Nature Reviews Drug Discovery, 2020, 19(3): 185–99.

[6] Depil S, Duchateau P, Grupp S A, et al. ‘Off-the-shelf’ allogeneic CAR T cells: development and challenges [J]. Nat Rev Drug Discov, 2020, 19(3): 185–99.

[7] Schmitz R, Fitch ZW, Schroder PM, et al. B cells in transplant tolerance and rejection: friends or foes? [J]. Transpl Int, 2020, 33(1): 30–40.

[8] Zachary AA, Leffell MS. HLA Mismatching Strategies for Solid Organ Transplantation -A Balancing Act [J]. Front Immunol, 2016, 7: 575.

[9] Wagner DL, Fritsche E, Pulsipher MA, et al. Immunogenicity of CAR T cells in cancer therapy [J]. Nat Rev Clin Oncol, 2021, 18(6): 379–93.

[10] Lei J, Ni Z, Zhang R. Universal CAR-T Cell Therapy for Cancer Treatment: Advances and Challenges [J]. Oncol Res, 2025, 33(11): 3347–73.

[11] Kagoya Y, Guo T, Yeung B, et al. Genetic Ablation of HLA Class I, Class II, and the T-cell Receptor Enables Allogeneic T Cells to Be Used for Adoptive T-cell Therapy [J]. Cancer Immunol Res, 2020, 8(7): 926–36.

[12] Ren J, Liu X, Fang C, et al. Multiplex Genome Editing to Generate Universal CAR T Cells Resistant to PD1 Inhibition [J]. Clinical Cancer Research, 2017, 23(9): 2255–66.

[13] Torikai H, Reik A, Soldner F, et al. Toward eliminating HLA class I expression to generate universal cells from allogeneic donors [J]. Blood, 2013, 122(8): 1341–9.

[14] Vivier E, Raulet DH, Moretta A, et al. Innate or adaptive immunity? The example of natural killer cells [J]. Science, 2011, 331(6013): 44–9.

[15] Long EO, Kim H S, Liu D, et al. Controlling natural killer cell responses: integration of signals for activation and inhibition [J]. Annu Rev Immunol, 2013, 31: 227–58.

[16] Parham P. MHC class I molecules and KIRs in human history, health and survival [J]. Nat Rev Immunol, 2005, 5(3): 201–14.

[17] Long EO. Negative signaling by inhibitory receptors: the NK cell paradigm [J]. Immunol Rev, 2008, 224: 70–84.

[18] Jo S, Das S, Williams A, et al. Endowing universal CAR T-cell with immune-evasive properties using TALEN-gene editing [J]. Nat Commun, 2022, 13(1): 3453.

[19] KäRre K. Natural killer cell recognition of missing self [J]. Nat Immunol, 2008, 9(5): 477–80.

[20] Gornalusse GG, Hirata RK, Funk SE, et al. HLA-E-expressing pluripotent stem cells escape allogeneic responses and lysis by NK cells [J]. Nat Biotechnol, 2017, 35(8): 765–72.

[21] Jandus C, Boligan K F, Chijioke O, et al. Interactions between Siglec-7/9 receptors and ligands influence NK cell-dependent tumor immunosurveillance [J]. J Clin Invest, 2014, 124(4): 1810–20.

[22] Ljunggren HG, KäRre K. In search of the ‘missing self’: MHC molecules and NK cell recognition [J]. Immunol Today, 1990, 11(7): 237–44.

[23] Parent AV, Faleo G, Chavez J, et al. Selective deletion of human leukocyte antigens protects stem cell-derived islets from immune rejection [J]. Cell Rep, 2021, 36(7): 109538.

[24] Guo Y, Xu B, Wu Z, et al. Mutant B2M-HLA-E and B2M-HLA-G fusion proteins protects universal chimeric antigen receptor-modified T cells from allogeneic NK cell-mediated lysis [J]. Eur J Immunol, 2021, 51(10): 2513–21.

[25] Rongvaux A, Willinger T, Martinek J, et al. Development and function of human innate immune cells in a humanized mouse model [J]. Nat Biotechnol, 2014, 32(4): 364–72.

[26] Walsh NC, Kenney LL, Jangalwe S, et al. Humanized Mouse Models of Clinical Disease [J]. Annu Rev Pathol, 2017, 12: 187–215.

[27] Nahmad A D, Reuveni E, Goldschmidt E, et al. Frequent aneuploidy in primary human T cells after CRISPR-Cas9 cleavage [J]. Nat Biotechnol, 2022, 40(12): 1807–13.

[28] Kosicki M, Tomberg K, Bradley A. Repair of double-strand breaks induced by CRISPR-Cas9 leads to large deletions and complex rearrangements [J]. Nat Biotechnol, 2018, 36(8): 765–71.

[29] Joyce JA, Fearon DT. T cell exclusion, immune privilege, and the tumor microenvironment [J]. Science, 2015, 348(6230): 74–80.

[30] Cherkassky L, Morello A, Villena-Vargas J, et al. Human CAR T cells with cell-intrinsic PD-1 checkpoint blockade resist tumor-mediated inhibition [J]. J Clin Invest, 2016, 126(8): 3130–44.

[31] Veglia F, Sanseviero E, Gabrilovich DI. Myeloid-derived suppressor cells in the era of increasing myeloid cell diversity [J]. Nat Rev Immunol, 2021, 21(8): 485–98.

[32] Grosser R, Cherkassky L, Chintala N, et al. Combination Immunotherapy with CAR T Cells and Checkpoint Blockade for the Treatment of Solid Tumors [J]. Cancer Cell, 2019, 36(5): 471–82.

